# Bioluminescence-based reporters for characterizing inhibitors and activators of human Sonic Hedgehog protein autoprocessing in live cells at high throughput

**DOI:** 10.1101/2022.06.27.497760

**Authors:** Daniel A Ciulla, Patricia Dranchak, John L Pezzullo, Rebecca A Mancusi, Alexandra Maria Psaras, Ganesha Rai, José-Luis Giner, James Inglese, Brian P Callahan

## Abstract

The Sonic hedgehog (SHh) precursor protein undergoes biosynthetic autoprocessing to cleave off and cholesterylate the SHh signaling ligand, a vital morphogen and oncogenic effector protein. Autoprocessing is self-catalyzed by SHhC, the SHh precursor’s enzymatic domain. Here we describe the development and validation of the first cellular reporter to monitor human SHhC autoprocessing non-invasively in high-throughput compatible plates. The assay couples intracellular SHhC autoprocessing to the extracellular secretion of the bioluminescent nanoluciferase enzyme. We developed a wild-type (WT) SHhC reporter line for evaluating potential autoprocessing inhibitors by concentration response-dependent suppression of extracellular bioluminescence. A conditional mutant SHhC (D46A) reporter line was developed for identifying potential autoprocessing activators by a concentration response-dependent gain of extracellular bioluminescence. The D46A mutation removes a conserved general base that is critical for the substrate activity of cholesterol. Inducibility of the D46A reporter was established using a synthetic sterol, 2-α carboxy cholestanol, designed to bypass the defect through intra-molecular general base catalysis. To facilitate direct nanoluciferase detection in the cell culture media of 1536-well plates, we designed a novel membrane-impermeable nanoluciferase substrate, CLZ-2P. This new reporter system offers a long-awaited resource for small molecule discovery for cancer and for developmental disorders where SHh ligand biosynthesis is dysregulated.

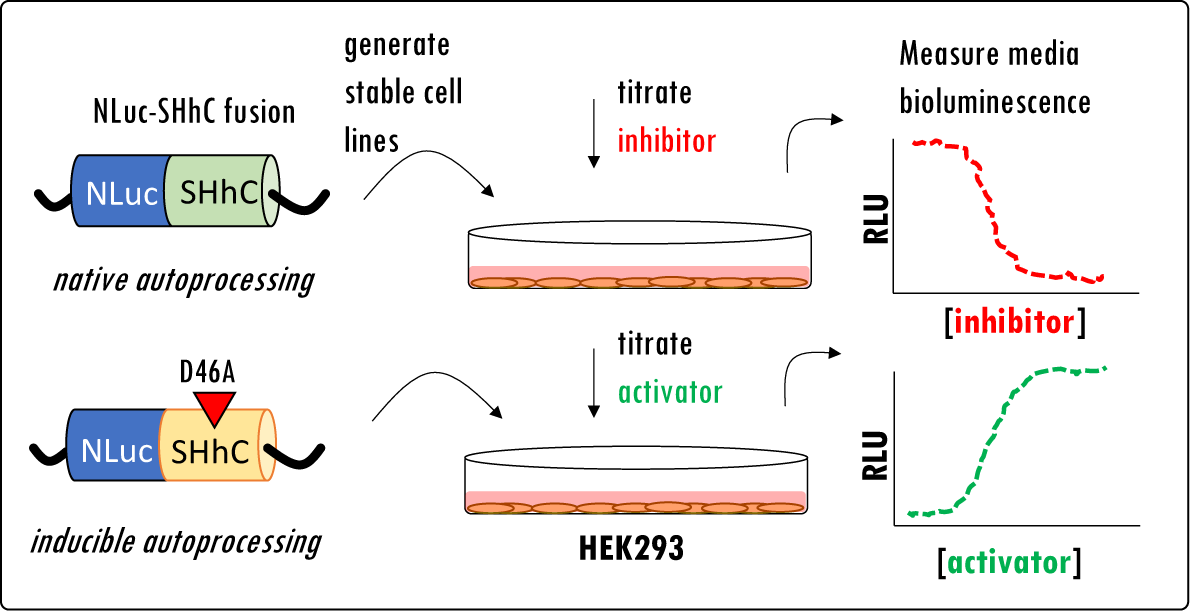

## INTRODUCTION

Sonic Hedgehog (SHh) ligands initiate vital cell signaling necessary for brain development and neural stem cell proliferation, while displaying oncogenic activity when dysregulated in sporadic tumors.^1-7^ The role of SHh signaling in health and disease, particularly in cancer, has prompted efforts to identify clinically useful antagonists of different pathway components.^8-12^ Many SHh signaling inhibitors target Smoothened (SMO), a cell surface receptor involved in SHh signal transduction^8^ ; however mutational escape from the anti-tumor effects of those SMO inhibitors presents a challenge to durable remission.^13-15^ Thus, new targets to suppress oncogenic SHh ligand signaling are of interest. In a different context, activating the SHh pathway holds promise for developmental disorders where the native function of SHh is compromised by diminished expression. Mutations in the human *SHh* gene are implicated in holoprosencephaly (HPE), a devasting congenital disorder affecting fetal brain development.^16^ Roughly half of the reported HPE-associated mutations in *SHh* map to the gene’s 3’ region,^5^ encoding a partially characterized autoprocessing domain called SHhC. In this context, small molecules that enhance SHh signaling by restoring SHhC function could find therapeutic application.

**Scheme 1.**
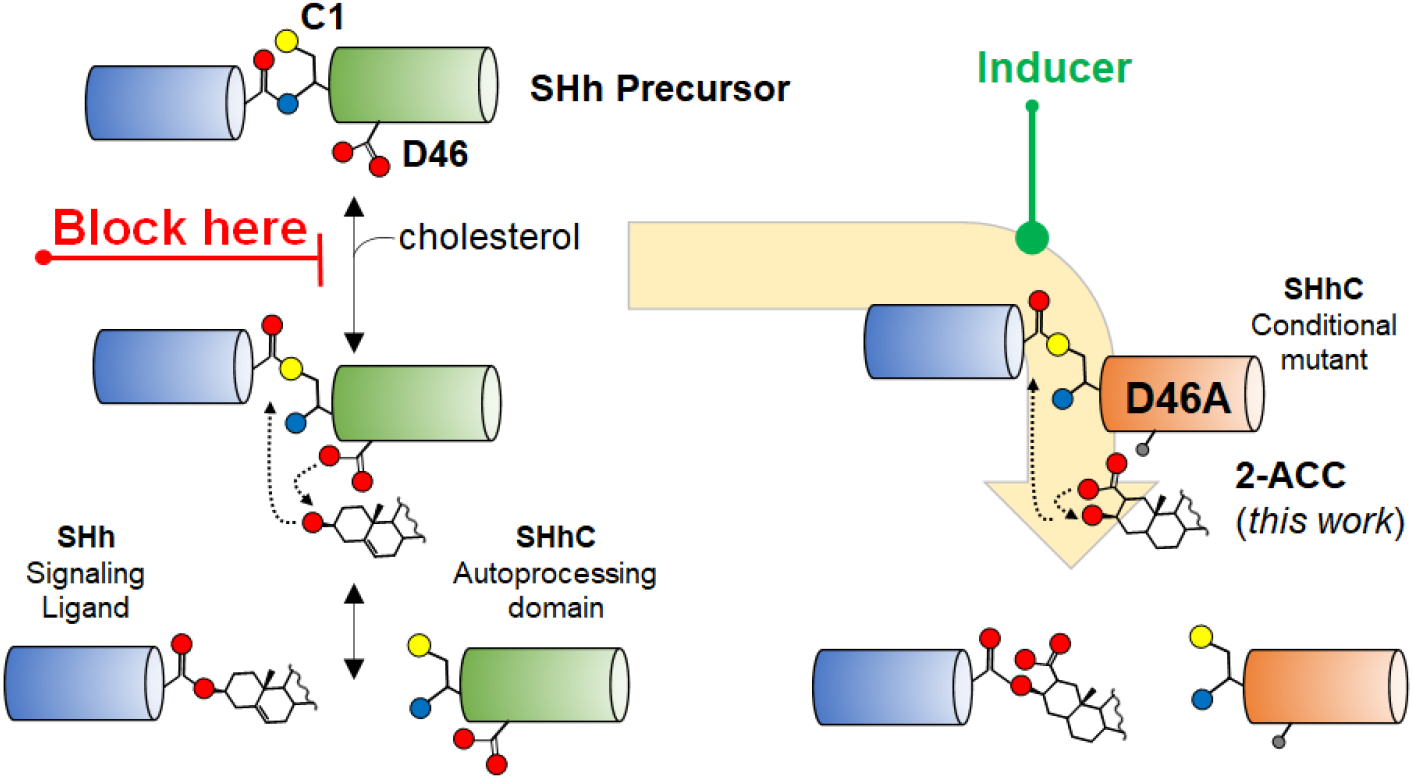
Native Hedgehog SHhC autoprocessing with substrate cholesterol and non-native, induced autoprocessing using chemical rescue approach. *Left*. In the native cholesterolysis pathway, SHhC (green) activates then cleaves off the adjacent SHh ligand (blue) using cholesterol as the terminal nucleophile. *Right* Defective autoprocessing by SHhC point mutant, D46A, is reactivated by engineered sterols, such as 2-α carboxy cholestanol (2-ACC) described here.

The autoprocessing SHhC domain is present in the precursor form of SHh where it self-catalyzes an essential biosynthetic step called cholesterolysis. Cholesterolysis occurs in the endoplasmic reticulum (ER) and results in the release and carboxyl-terminal cholesterylation of the adjacent SHh ligand domain **(Scheme 1, *left***).^17-19^ The native reaction begins with a backbone peptide bond rearrangement at the junction between the SHh ligand and the SHhC domain. This rearrangement activates the terminal Gly residue of the SHh ligand as a thioester at the Cys (C1) side chain of SHhC. Mechanistic overlap exists here with self-splicing inteins.^20, 21^ In a step unique to Hh family proteins, SHhC then catalyzes nucleophilic attack at the internal thioester by a cholesterol molecule. A conserved aspartate residue of SHhC, Asp46 (D46), is the putative general base for the attacking C3-OH group of cholesterol.^22^ Cleavage releases cholesterylated SHh ligand for N-palmitoylation, Golgi transport, extracellular secretion and downstream cell/cell signaling. Mutations that deactivate SHhC result in ER retention of the SHh precursor and lead to its eventual removal through Endoplasmic Reticulum Associated Degradation (ERAD).^23-25^ Engineered sterols have the potential to circumvent catalytic defects in SHhC and restore SHh ligand biosynthesis in some autoprocessing mutants (**Scheme 1, *right***).(Ref ^26^ and this work)

Here we describe and validate the first homogenous live cell autoprocessing assay to screen for inhibitors and activators of human SHhC function. Chemical activators of SHhC could mitigate the impact of autoprocessing mutations, diminishing HPE pathology; antagonists of SHhC provide a novel means to block oncogenic biosynthesis of the Sonic Hh ligand.^13, 27^ Our method couples intracellular SHhC autoprocessing to the extracellular secretion of the ATP-independent nanoluciferase (NLuc) enzyme.^28^ A Sonic Hedgehog wild-type (WT) autoprocessing reporter line is established in HEK293 cells for evaluating SHhC inhibitors by a concentration-dependent loss of extracellular NLuc bioluminescence. We also describe and validate an inducible autoprocessing mutant (D46A) for identifying SHhC activators by the gain of extracellular NLuc bioluminescence. Direct NLuc detection in the media of 1536-well cell culture plates is facilitated by using methylphosphonic acid coelenterazine, a novel cell impermeable anionic substrate analog for NLuc.

## 2. RESULTS

### 2.1 Reporter Strategy

Autoprocessing activity of precursor Sonic Hh protein appears to reside exclusively in SHhC. The SHh ligand is a spectator to the transformation and this arrangement allows for the substitution of the SHh ligand with heterologous polypeptides. Chimeric, functional SHh reporter constructs have been described where green fluorescent protein is fused to SHhC, and more recently, where Halo-Tag is fused to SHhC.^29, 30^ We substituted the SHh ligand with HA-tagged nanoluciferase^28^ (HA-NLuc) with a view toward an HTS-compatible assay system.

Three related HA-NLuc-SHhC fusion constructs were prepared (**Figure 1 A**). First, a wild-type (**WT**) HA-NLuc-SHhC was designed for assessment of native SHhC activity and to characterize potential autoprocessing inhibitors. Autoprocessing of the **WT** reporter construct is expected to cleave off and cholesterylate HA-NLuc using cellular cholesterol in the ER. Cholesterylated HA-NLuc would then transit through the secretory pathway for extracellular release. NLuc activity present in the culture media thereby reports cellular SHhC autoprocessing. The N-terminal HA tag of NLuc enables the analysis of SHhC autoprocessing by Western blot. The **WT** reporter construct is intended for evaluating autoprocessing inhibitors by a decrease in extracellular NLuc and by an accumulation of intracellular **WT** precursor.

**FIGURE 1.**
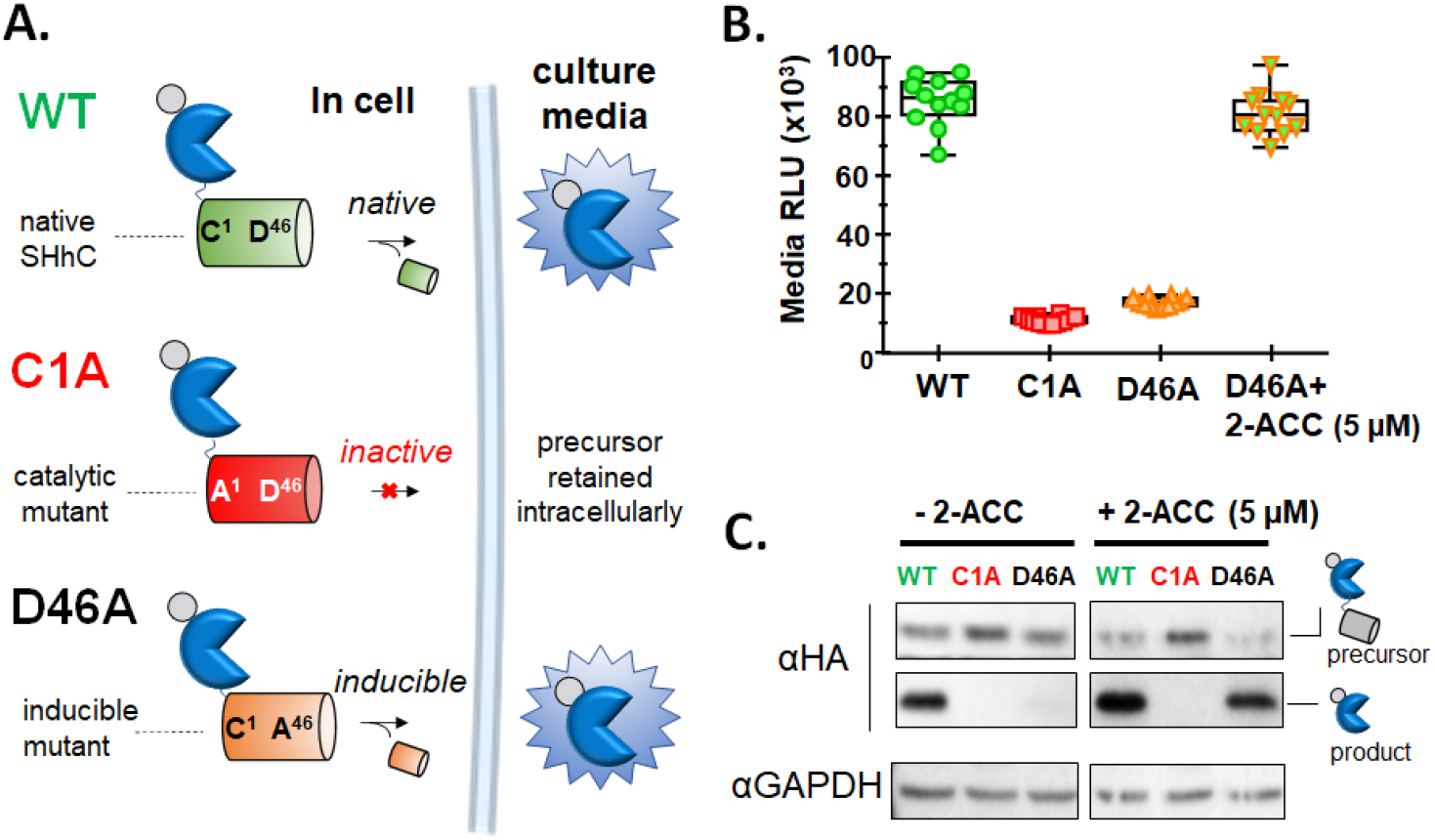
Monitoring autoprocessing in live cells using HA-tagged nanoluciferase (HA-NLuc) as a secreted, proxy reporter of SHhC activity. A. Autoprocessing reporter constructs. **WT**, constitutively active. autoprocessing using cellular cholesterol; **C1A**, irreversibly inactive autoprocessing mutant; **D46A**. conditional mutant with inducible autoprocessing activity. Reporter constructs carry an N-terminal HA epitope tag (grey) for Western analysis and the NLuc (blue) enzyme for non-invasive measurements of autoprocessing by media bioluminescence. **B**. Media bioluminescence reflects intracellular SHhC autoprocessing. Functional behavior of the **WT, C1A**, and **D46A** reporters was evaluated after transient transfection in HEK293 cells (n=12). Inducibility of the D46A reporter was assayed by comparison of +/- the 2-ACC activator (5 μM). C. Western blot for **WT, C1A** and **D46A** precursor and product polypeptides using the N-terminal HA epitope. Cell extract was analyzed for the three reporter constructs in the absence (left) and presence (right) of 2-ACC (5 μM). GAPDH served as a loading control.

We prepared a control construct, **C1A**, to mimic 100% inhibition of SHhC autoprocessing. This reporter fuses HA-NLuc to a **C1A** mutant of SHhC. The Cys>Ala mutation at position 1 of SHhC removes the critical nucleophilic thiol group required for thioester formation, thereby blocking SHhC enzymatic function. The precursor form of the **C1A** reporter is expected to remain largely in the cellular fraction where it would be subject to degradation by ERAD. Compounds that act specifically toward SHhC autoprocessing are not expected to influence bioluminescence from the catalytically inactive **C1A** or alter the reporter protein’s steady-state expression when analyzed by Western blot.

In the third construct, we fused HA-NLuc to a **D46A** point mutant of SHhC for an autoprocessing reporter with inducible activity. The **D46A** construct is intended for evaluating small molecules as potential autoprocessing activators. Earlier studies on *Drosophila* Hh autoprocessing indicated that **D46A** mutants of HhC can form internal thioester; however, without the conserved active-site D46 carboxylate group, transesterification to substrate cholesterol is prevented.^22^ The corresponding **D46A** mutation in human SHhC was expected to behave in a similar manner, blocking native cholesterolysis in cells. Based on our experiments with *Drosophila* HhC, we hypothesized that internal thioester generated by **D46A** could be resolved with rationally designed analogs such as 3-HPC (3-β hydroperoxycholestane) that can bypass the mutant’s catalytic defect.^26^ For the *in vitro* cell culture work here, we prepared a more chemically stable hyper-nucleophilic analog, 2-α carboxy cholestanol (2-ACC). The carboxylate group vicinal to the attacking C3 hydroxyl group in 2-ACC was designed to functionally replace the missing carboxylate in the **D46A** mutant. (**Scheme 1**, *right*; and **Figure 2**). Below, we demonstrate that 2-ACC serves as an intracellular inducer of **D46A**.

**Figure 2.**
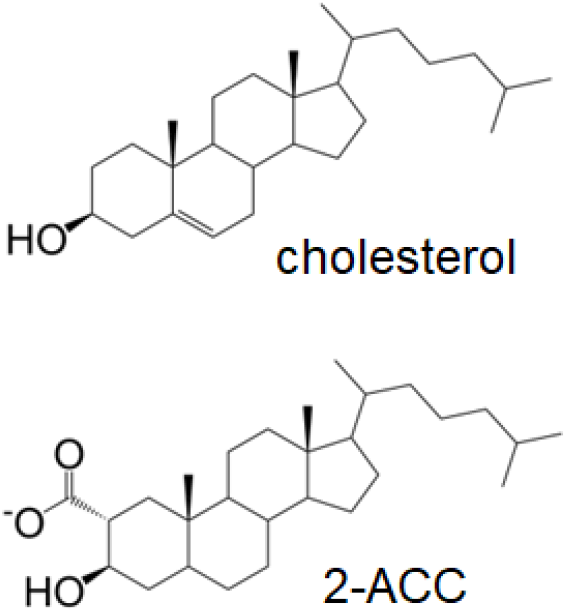
Substrate cholesterol for **WT** reporter autoprocessing and synthetic activator substrate, 2-ACC, for inducing autoprocessing by the **D46A** reporter.

### 2.2 Nanoluciferase (NLuc) as a surrogate SHhN ligand and extracellular bioluminescent reporter

Initial characterization of the SHhC autoprocessing reporters was carried out by transient transfection using HEK293 cells. HEK293 have been used before for cellular studies of human SHhC autoprocessing.^31, 32^ The **WT, C1A** and **D46A** chimeric precursors were cloned into the pDisplay (Invitrogen) mammalian expression vector in frame with the vector’s N-terminal Ig κ-chain secretion sequence and HA epitope tag. Following transfection in HEK293, SHhC activity was assayed by NLuc bioluminescence in the culture media and by Western blot of the cell extract.

**Figure 1B** summarizes media bioluminescence using the NLuc NanoGlo reagent (Promega) following transient transfections with the three reporter plasmids. The first column in **Figure 1B** (green circles) shows readings from the **WT** samples. The mean bioluminescence value for the **WT** transfected cells was consistently the highest of the four conditions tested here, in accord with fully active SHhC autoprocessing. The catalytic mutant, **C1A** (**Figure 1B**, red squares) by contrast had the lowest output, reduced from **WT** by 6.5-fold, consistent with defective autoprocessing and intracellular precursor retention.^23^

The last two columns in **Figure 1B** show the media bioluminescence from HEK293 cells transfected with the **D46A** reporter plasmid (–) and (+) 2-ACC (5 μM). In samples lacking 2-ACC, the **D46A** samples produced media bioluminescence that is on par with the **C1A** reporter. This correspondence is consistent with the inability of **D46A** to autoprocess using cellular cholesterol. The last column in **Figure 1B** showing **D46A** (+) 2-ACC indicates a clear gain of media bioluminescence (4.6-fold) relative to **D46A** without 2-ACC. The enhanced signal is exciting and supports the induction of autoprocessing in HEK293 cells by added 2-ACC. Examples like this where “chemical rescue” of mutant enzymes takes place in the cell are rare.^33-35^ Induced cellular autoprocessing of **D46A** by 2-ACC is further supported by Western blot analysis, described next. In the SI material, we also report that the corresponding D46A mutant of *Drosophila* HhC has its defective autoprocessing activity restored by 2-ACC (**Supplementary Figure 1**). Bioluminescence was not appreciably enhanced when 2-ACC was added to cells transfected with the **C1A** or **WT** reporters, establishing the specificity of this effect. Overall, the results of the transient transfection experiments in **Figure 1B** provide proof-of-concept for the **WT, C1A** and **D46A** reporters.

To complement the comparison based on NLuc bioluminescence, we probed the corresponding cellular fractions of **WT, C1A** and **D46A** by Western blot using the construct’s HA epitope tag (**Figure 1C**). Lysate samples were denatured, separated by SDS-PAGE and probed with a monoclonal HA-HRP antibody (Sigma) to identify precursor protein, HA-NLuc-SHhC (52 kDa) and autoprocessing product, HA-NLuc (24.4 kDa). Analysis of samples with the **WT** reporter confirmed SHhC autoprocessing activity, with precursor and product in a ratio of ∼1 to 10. Lysate from the **C1A** samples showed signal for precursor only, consistent with deactivated SHhC autoprocessing. In **D46A**, induction of autoprocessing was evident by comparison of samples (-)/(+) 2-ACC. In the absence of 2-ACC, the **D46A** precursor is expressed without appreciable generation of autoprocessing product. By contrast, signal for the **D46A** precursor decreases in the (+) 2-ACC sample and the HA-NLuc product signal accumulates, consistent with restored SHhC autoprocessing. The induction was shown to be specific to **D46A** (**Figure 1C**, compare - /+ 2-ACC sample sets for **D46A** with **WT** and **C1A**).

### 2.3 Miniaturizing the NLuc-SHhC autoprocessing reporter system for 1536 well plates

With a view toward applications to high throughput chemical screening for inhibitors and activators^36^ of SHhC we sought to streamline assay setup by first preparing stable **WT** and **D46A** reporter cell lines. CRISPR/Cas9 targeted gene insertion^37^ was used to integrate the two reporter constructs into HEK293 cells (*Materials and Methods*).

In line with the transient transfection results above, the stable **WT** line showed constitutive autoprocessing with strong bioluminescence output in the culture media. We found that treatment of the **WT** line with brefeldin A, a protein secretion blocker that disrupts ER to Golgi transport, attenuated media bioluminescence, supporting the notion that Nluc activity in the media reflects secretory pathway transport rather than non-specific cell leakage (**Figure 3A**). For additional validation, we tested whether the **WT** reporter would recapitulate the ERAD/proteasome sensitivity observed with native human Sonic Hh precursor. ^23^ Studies by Salic *et al* suggest that the human SHh precursor in the ER is subjected to a dynamic partitioning between native autoprocessing and ERAD-mediated proteasome degradation. We monitored **WT** precursor levels by Western blot following various treatments. We first treated the **WT** line with the protein synthesis inhibitor cycloheximide (CHX). With CHX, we observed a time dependent precursor loss (**Fig. 3B**, CHX, top row of Western blot), which could be attributed to native autoprocessing as well as to ERAD. For comparison, in the untreated (DMSO only) samples, the HA-NLuc-SHhC precursor appears to be at a steady state (**Fig. 3B**, no treatment). To separate the influence of native autoprocessing from ERAD degradation, the **WT** line was treated initially with the proteasome inhibitor MG132. Inhibiting the proteasome resulted in a ∼70% increase in the precursor signal. Precursor accumulation is consistent with diminished ERAD (**Fig 3B**, MG132). Next, the **WT** cells were treated with a combination of CHX and MG132. Here we observed a more gradual consumption of precursor (**Fig 3B**, MG132+CHX). Under this final condition, precursor levels should predominately reflect SHhC autoprocessing with endogenous cholesterol.^23^ Overall, the pattern of accumulation and depletion of the **WT** precursor protein in **Figure 3B** agrees with the dynamic partitioning between ERAD-mediated destruction and SHhC autoprocessing observed with native SHh precursor.

**FIGURE 3.**
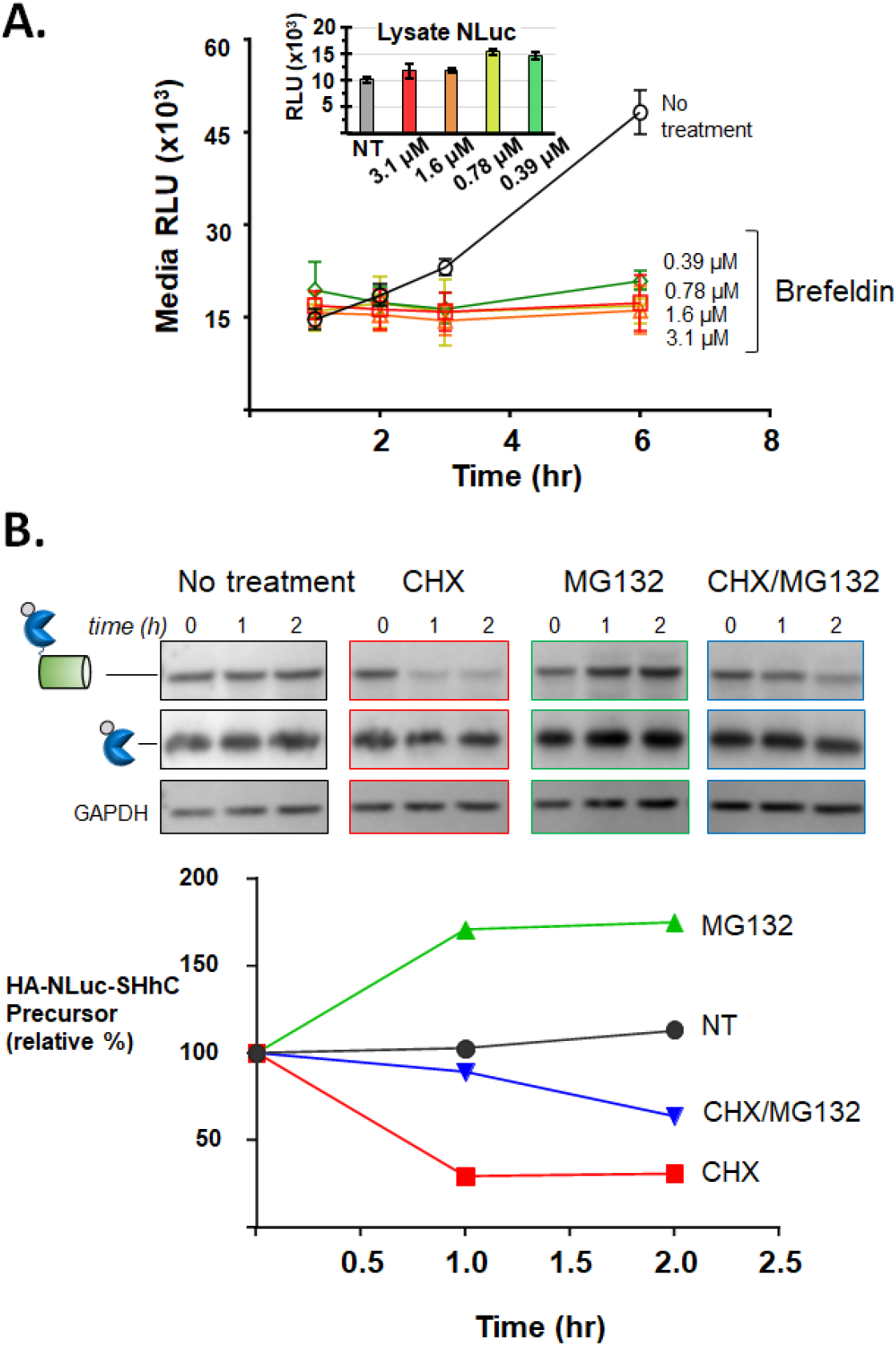
Validation of the WT line as a reporter for native SHhC autoprocessing. **A**. Extracellular secretion of NLuc by **WT** reporter blocked by Brefeldin A. Media bioluminescence was measured 1, 2, 3, and 6 hours after addition of the indicated concentrations of Brefeldin A. (inset) Bioluminescence accumulates in cell lysate fraction after 6 hr Brefeldin A. **B**. Intracellular **WT** reporter protein response to cycloheximide (CHX) and MG132. Upper. Western Blot analysis. Control samples treated with DMSO control; 25 μg/ml cycloheximide; 50 μM MG132; and co-treated with 25 μg/ml CHX and 50 μM MG132. *Lower*. Plot of **WT** precursor abundance using Western blot images shown based on analysis with Image J.

The stable HEK293 **D46A** reporter line displayed specific, inducible autoprocessing activity as measured by extracellular NLuc activity and in accord with the transient transfection experiments. In the absence of 2-ACC, the **D46A** reporter line again displayed low media bioluminescence, consistent with intracellular retention. As shown in **Figure 4A**, there was a gain of media bioluminescence with added 2-ACC that was concentration-dependent and saturable over an 11-pt titration, from 20 nM to 20 μM in 2-ACC. Analysis of the concentration-response data by non-linear regression yielded an EC_50_ value of 1.6 μM for 2-ACC. As an additional control, we added cholesterol to the **D46A** line at a concentration equivalent to 2-ACC (5 μM); no increase in media bioluminescence was apparent (**Figure 4B**). The hyper-nucleophilic 3-HPC failed to produce measurable induction of cellular autoprocessing by **D46A**, likely a consequence of the hydroperoxide’s instability.

**FIGURE 4.**
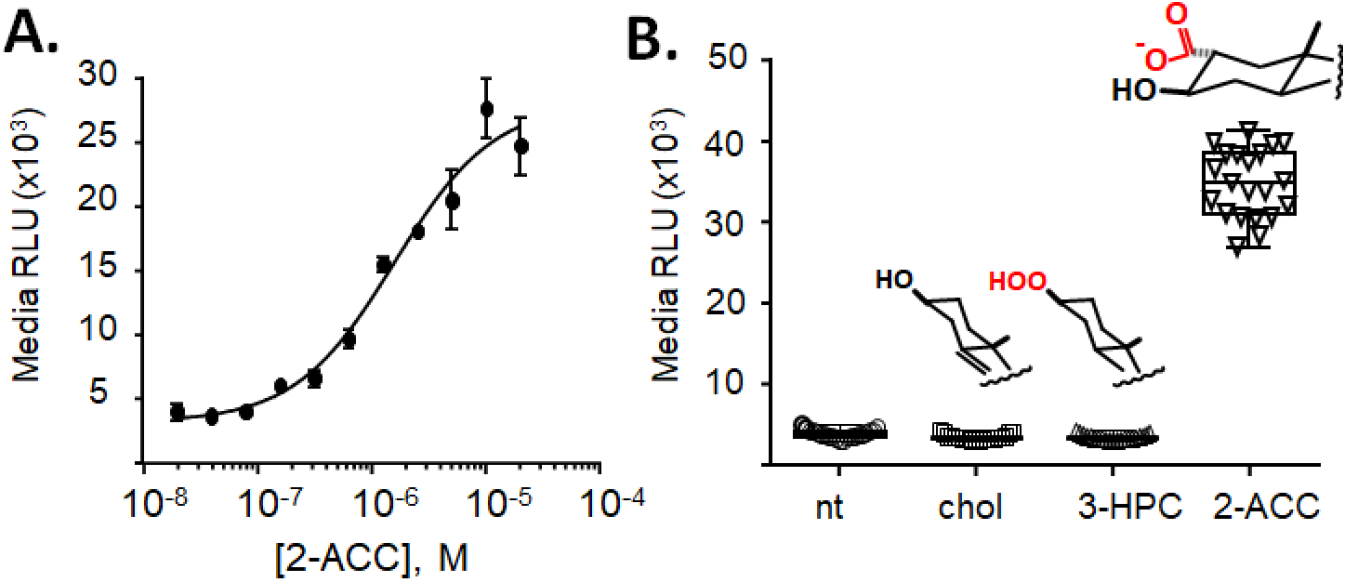
Validation of the D46A line as a chemically inducible SHhC autoprocessing reporter. **A**. Concentration-response activation curve for 2-ACC based on media NLuc activity from **D46A** (n=4). EC_50_ = 1.6 μM. **B**. Induced autoprocessing with **D46A** is 2-ACC specific. Endogenous cellular cholesterol (no treatment), added cholesterol (5 μM), added 3-HPC (5 μM) compared with added 2-ACC (5 μM) as inducers (n=21).

Next, we focused on optimizing and miniaturizing the **WT**/**D46A** reporter system to the 1536-well format. To streamline the assay, we designed and synthesized a cell-impermeable coelenterazine analog (CLZ-2P). Traditional imidazopyrazinone nanoluciferase substrates such as coelenterazine (CLZ) and furimazine (NanoGlo) are membrane permeable and therefore require careful media transfer from the cell culture plate to an assay plate to separate signal from intra- and extracellular NLuc.^38^ Intracellular NLuc could include product *and* precursor forms of the reporter protein used here, confounding analysis of autoprocessing activity; whereas extracellular NLuc is expected to reflect autoprocessing product only. CLZ-2P, incorporating an anionic methylphosphonic acid at the 2-(4-hydroxybenzyl) moiety of CLZ was devised to eliminate the media transfer step for measurement of media bioluminescence (**Figure 5A**). The C-P phosphonyl bond in CLZ-2P was incorporated to stabilize the anionic group against P-O cleaving phosphatases, similar to the previously described analog described by Lindberg et al ^39^, but synthetically more tractable for quantities needed in HTS applications. The data in **Figure 5B** supports CLZ-2P as an alternative NLuc substrate that is suited for mix-and-read assays of the **WT** and **D46A** in 1536 well plates.

**FIGURE 5.**
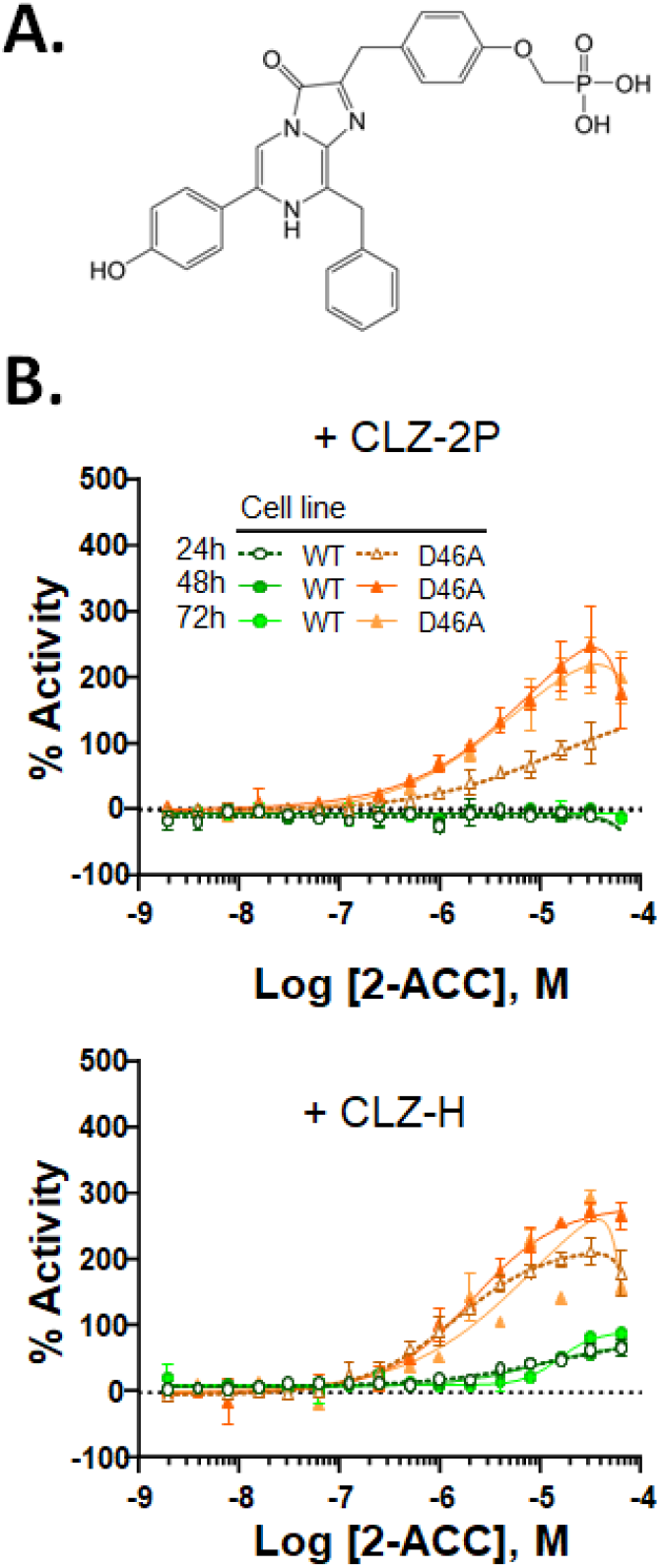
**A**. Anionic coelenterazine, CLZ-2R **B**. Validation of CLZ-2P for mix-and-read extracellular NLuc bioluminescence in 1536-well plates. Samples of WT and D46A with CLZ-2P (top) or CLZ-H (bottom) were analyzed as a function of increasing 2-ACC. Data were row-wise normalized to DMSO neutral control at each time point. Error bars represent standard deviation of the average of 2 replicate wells.

Analysis of the **WT** and **D46A** reporter lines in the 1536-well plate format was conducted hereafter using CLZ-2P. We observed a relatively low coefficient of variation of 4-5% between 32 intra-plate replicates for both cell lines (**Supplementary Table 1**), demonstrating well-to-well reproducibility in this assay format. Both reporter lines yielded a suitable Z’ factor, with **WT** = 0.83 and **D46A** = 0.64, using cycloheximide and 2-ACC, respectively.^36^ In **Figure 6 A-B**, the concentration-response curves are shown for **D46A** and **WT** treated with 2-ACC and CHX. In agreement with the experiments above, activation by 2-ACC is specific to **D46A**, with an EC_50_ of ∼5 μM and ∼100% activation at 28.75 μM 2-ACC. Extracellular secretion of NLuc by **WT** was also shown to be sensitive to CHX treatment, with maximum inhibition of NLuc activity in the media at 57.5 μM CHX. As an additional test, cells were treated with the membrane-disrupting digitonin toxin and cytotoxicity was measured with CellTiter-Glo assay reagent. **D46A** and **WT** responded near identically, with maximum lethal digitonin concentration of 115 μM, Z’ factor:0.87 (**Figure 6C**). Taken together, the robust performance of the two stable lines in HTS-compatible plates, along with the efficacy of CLZ-2P as an NLuc substrate for media bioluminescence measurements, encourage the application of this novel reporter system to identify the first chemical probes of SHhC autoprocessing.

**Figure 6.**
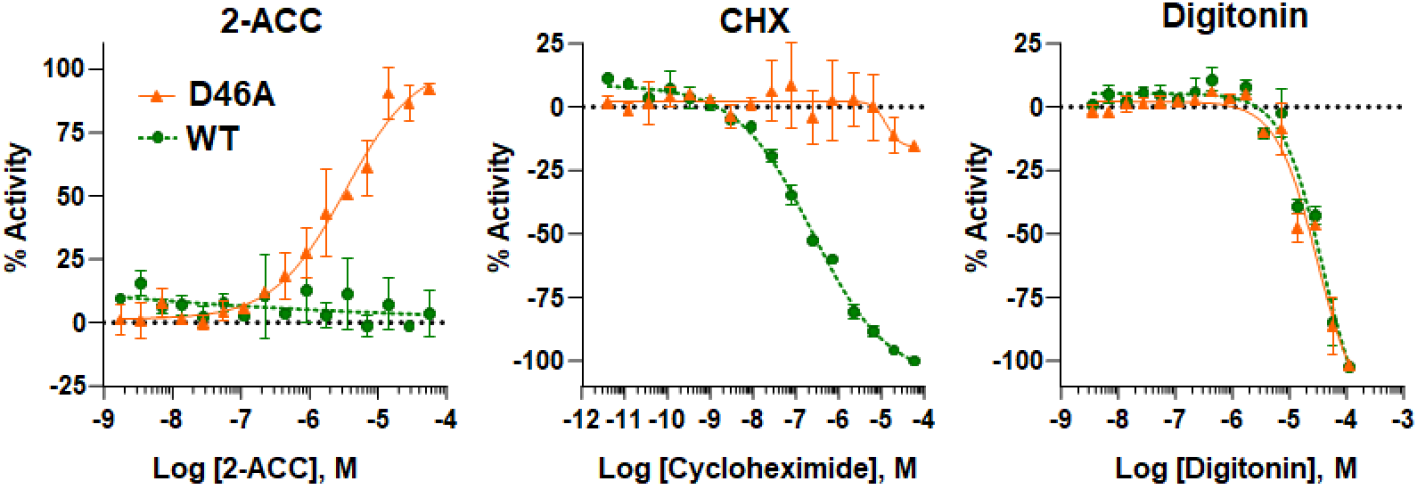
Concentration-response curves for WT and D46A in 1536-well plates. Graphs for 2-ACC (left) and cycloheximide (middle) are based on NLuc bioluminescence using CLZ-2P. Digitonin experiments are based on cytotoxicity, measured by the CellTiter-Glo reagent. Curves were fit in GraphPad Prism and error bars represent the standard deviation of 2 replicate wells for each concentration across the respective assay plates. n=2.

## DISCUSSION

Early studies on chicken SHh protein demonstrated “proteolytic processing” into two SHh products of approximately 19 kDa (SHh ligand) and 27 kDa (SHhC).^17^ Work carried out around the same time with mouse SHh reported similar behavior.^40^ Both studies involved transient expression of the full-length SHh precursor in cultured eukaryotic cells followed by Western blotting of cell lysate. Gel-based assays remain the standard approach to monitor cellular SHh autoprocessing despite the method’s idiosyncrasies, expense and low throughput. Cell-free and gel-free autoprocessing assays using recombinant protein, which we recently devised for kinetic studies of *Drosophila* Hh autoprocessing,^41, 42^ have so far resisted application to the human Hh homologs. In our hands, only sparing amounts of bacterially expressed human SHhC can be recovered, and that material seems to lack autoprocessing activity. Requirements for N-linked glycosylation^24^ and reductive activation^23^ of SHhC in the mammalian ER offer a possible explanation.

The *in vitro* live cell SHhC activity assay reported here represents an important resource for future studies of human Hh protein autoprocessing. With NLuc as the surrogate SHh ligand, intracellular SHhC autoprocessing is coupled to the secretion of a thermostable, ATP-independent bioluminescent reporter. Measurement of NLuc enzymatic activity is carried out non-invasively in a semi-quantitative manner in culture media. We show that bioluminescent output is sufficiently robust for miniaturization to 1536 well plates. Inhibitors of SHhC autoprocessing are expected to suppress NLuc secretion from cells expressing the **WT** reporter, while activators of SHhC autoprocessing are identified by enhanced NLuc secretion from the **D46A** reporter. The development of CLZ-2P as an impermeable substrate for NLuc increases assay throughput. With CLZ-2P, measurements of secreted NLuc are completed as a simple mix-and-read step in the presence of live cells.

The **WT** and **D46A** reporter cell lines are poised for quantitative high-throughput screening (qHTS) ^43^ to support the search for chemical probes of this key initiating step in the Sonic Hh signaling pathway. Autoprocessing events like Hh protein cholesterolysis that are single turnover and partly unimolecular represent challenging but not untenable targets for small molecules.^42, 44-47^ Antagonists of SHhC autoprocessing are sought to inhibit Sonic Hh biosynthesis, blocking the ligand from oncogenic signaling in sporadic tumors. In developmental disorders like HPE where SHh ligand biosynthesis is compromised by congenital mutation, activators of SHhC could serve as chemical chaperones to restore SHh signaling.

## Acknowledgements

This research was supported (in part) by the Intramural Research Program of the National Center for Advancing Translational Sciences (NCATS), National Institutes of Health (NIH) under project 1ZIATR000053 (J.I.), by the National Cancer Institute, Grant R01 CA206592 (BPC) and by National Institutes of General Medical Sciences, Grant R15GM143714 (JLG)

## MATERIALS and METHODS

### Biochemicals and cell biology reagents

DNA restriction enzymes, T4 DNA ligase and chemically competent DH5 alpha cells (NEB); Alt-R CRISPR-Cas9 crRNA, Cas9 expression plasmid, and PCR oligos (IDT). HA-HRP primary antibody; GAPDH-HRP primary antibody (Santa Cruz SC-365062); Hyclone 0.25% Trypsin-EDTA (GE Life Sciences); penicillin/streptomycin, EMEM, FBS (Corning); G418, OptiMEM I (Gibco); digitonin, MG132, cycloheximide (Millipore Sigma); HEK293 cells (ATCC, provided by Tracy Brooks, SUNY Binghamton). Plates: 96 Well Black Polystyrene Microplate (Corning 3650), 12 Well Nunclon Delta Surface (Thermo Scientific), 96 Well Tissue Culture Plate (VWR 734-2327), 50 mL Tissue Culture Flask Blue Vented Cap (Corning 353108), white solid bottom TC treated 1536 well plate (Greiner Bio-One 789173-F).

### Reporter Constructs

Chimeric SHh precursor encoding HA-tagged NLuc fused to native and mutant forms of human Sonic HhC autoprocessing domain were assembled in pDisplay (Invitrogen). Fragments encoding Sonic HhC were inserted into pDisplay using PstI and XhoI; codon optimized NLuc was inserted using the vector’s BglII and PstI sites. An oligo cassette for CRISPR-Cas9 genomic integration was introduced into the vector at a unique SnaBI site. Plasmids were purified for transfection with the ZymoPURE II plasmid midiprep kit (Zymoresearch). We thank Dr. Erich Roessler (NIH) for providing a Sonic Hh cDNA plasmid.

### Transient Transfection

HEK293 cells were seeded into a 12-well plate at 100,000 cell/mL in complete EMEM supplemented with 10% FBS and 0.1 % Pen/Strep. When cells reached 70% confluency, media was replaced with reduced serum OptiMEM I containing lipofectamine-2000 with 1.6 μg/well reporter plasmid. After a one-hour transfection, media was replaced with complete EMEM and the cells were incubated overnight. Cells and media were then collected for bioluminescence assays and Western blot analysis.

### HEK293 Stable Cell Line Generation using CRISPR/CAS9

We followed the general method outlined by He *et al*. to generate **WT** and **D46A** reporter lines.^37^ HEK293 cells were cultured at 37°C 5% CO_2_ in EMEM supplemented with 10% FBS and 0.1% Pen/Strep. Cells were grown to 70% confluency in a 12-well plate. The media was replaced with reduced serum OptiMEM I containing lipofectamine-2000 (Invitrogen), the pDisplay reporter plasmid (0.6 μg), Cas9 expression plasmid (0.6 μg), and short guide RNA (0.4 μg). After one hour, the transfection media was replaced with complete media. Following an overnight recovery in complete EMEM, the media was exchanged with complete EMEM containing G418 antibiotic (400 μg/ml). Media was then replaced every 2-3 days and cells were passaged weekly. Bioluminescence measurements were made periodically to confirm expression of the reporter constructs. After four weeks under G418 selection, cells were diluted into 96-well plates at single cell per well density and then expanded into stable cell lines.

### Cell lysis and Western blot analysis

HEK293 cells transfected with **WT, C1A** or **D46A** reporter plasmids were lysed after media aspiration by 250 μl ice-cold RIPA buffer (Thermo Scientific). After agitation by orbital shaking for 20 minutes at room temperature, the lysate suspension was collected into microcentrifuge tubes, treated with DNAse I (12 ng/μl) on ice for ten minutes, followed by centrifugation at 18,000 g for 5 minutes at 4°C. The supernatant was transferred to a clean tube and used immediately or stored at -80 °C. For Western analysis, aliquots of the soluble lysate were denatured by boiling at 95°C for 5 minutes in SDS-PAGE load buffer (final conc. Bromophenol blue 0.1%, glycerol 10%, SDS 2%, DTT 100 mM, Tris-HCl 50 mM). Denatured proteins were resolved by 12% SDS-PAGE. Following electrophoresis, the gel was submerged for 20 minutes in transfer buffer (Tris 2.5 mM, glycine 19 mM, 20% methanol, pH 8.3) then electroblotted (20 V) overnight at 4°C onto a 0.2 μm PVDF membrane (Amersham Hybond). Following transfer, the membrane was blocked with 5% BLOT-Quick at room temperature with gentle shaking for one hour. After rinses with TBST buffer (Tris 2 mM, NaCl 15 mM, Tween-20 0.1%), a solution of TBST/5% BLOT-quick with αHA-HRP (1/500) or αGAPDH-HRP (1/1000) was added. After gentle shaking at 4 °C, overnight, the membrane was rinsed 3x with TBST buffer, followed by a 15-minute wash, and then three five-minute washes with TBST buffer. Visualization of HRP was accomplished by colorimetric detection (Femto-Chromo HRP kit, G-Biosciences).

### Extracellular NLuc Assay for induced autoprocessing in the D46A line

Stable **D46A** cells were seeded into 96-well plates at a density of 2×10^4^ cells/well (total volume 100 μl) and grown to ∼70% confluency over 72 hours. Seeding media contained 2-ACC, cholesterol, or 3-HPC, titrated over an 11-point 1:2 dilution from 20 μM to 20 nM. At least four intra-plate replicates were included for each concentration of sterol. We assayed 10 μl of the media for bioluminescence using the NanoGlo detection reagent (furimazine). We first prepared a NLuc reaction stock solution by combining 980 μl NanoGlo buffer with 20 μl of furimazine substrate. This solution was incubated in the dark at room temperature for five minutes. During this period, we prepared the bioluminescence assay plate. Each sample well in the assay plate (Corning 3650) contained 90 μl of distilled water and 10 μl media from the cell culture plate. To initiate bioluminescence, 10 μl of the NanoGlo reaction solution was added to each sample well in the assay plate, incubated with gentle shaking for two minutes, then assayed for bioluminescence using a Biotek Synergy H1 plate reader with gain equal to 150.

### Assay miniaturization and NLuc substrate optimization

HEK293 cells expressing wildtype HA-NLuc-SHhC (**WT**) and HA-NLuc-SHhC (**D46A**) were grown in Eagle’s Minimum Essential Medium (ATCC) supplemented with 10% FBS (HyClone), 100 U/ml penicillin, and 100 μg/ml streptomycin (Gibco) in the absence of selection antibiotics. Cells were trypsinized from culture flasks, counted, diluted in growth media and plated at 1000, 1500, or 2000 cells per 4 μl per well in respective columns of white, solid bottom TC treated 1536-well plates (Greiner Bio-One) with a multidrop combi dispenser (ThermoFisher). Control compound digitonin was prepared at 20 mM stock concentration in DMSO and diluted in a 16-pt, 1:2 dilution, while MG132 and cycloheximide were prepared at a stock concentration of 10 mM in DMSO and titrated in 16-pt, 1:3 dilution. Each compound titration was transferred in duplicate to respective columns of assay plates along with replicate columns of DMSO vehicle control to seeded cells at 25 nL per well with a Mosquito liquid handler (SPTLabTech) immediately after cells were plated. Cells were covered with weighted metal lids with gas exchange pores (Wako) and incubated at 37°C, 5% CO_2_ and 95% humidity. After 24 hr 1 μl media was manually transferred from each well of one assay plate to a recipient white, solid bottom TC treated 1536-well plate with 3 μl 1x PBS predispensed in each well. NanoGlo luciferase assay substrate was prepared at a 1:100 dilution according to manufacturer’s protocol and 3 μl NanoGlo substrate was added to each well of media transfer plate and cell source plate with a BioRAPTR FRD (Beckman Coulter). Nanoluciferase luminescence for each plate was read on a ViewLux plate reader (PerkinElmer) after 5 min. Coelenterazine-H substrate was prepared as a 2 mM stock solution in 100% ethanol and coelenterazine-2P was diluted to a 2 mM stock solution in DMSO. Both substrates were prepared in a non-lytic PBS assay buffer (300 mM sodium ascorbate, 5 mM NaCl) at a working dilution of 20 μM. Coelenterazine substrates were added at 4 μl/well to one cellular assay plate each with a BioRAPTR FRD and nanoluciferase luminescence was read on a ViewLux plate reader within 2-5 min.

### qHTS 2-ACC Time Course Concentration-response Validation

Stable HEK293 **WT** and **D46A** cells were grown and plated at 1.5×10^3^ cells/well (4 μl) in respective columns of six replicate white, solid bottom TC treated 1536-well plates as before. 2-α carboxy cholestanol was diluted in a 16-pt, 1:2 dilution starting at a high concentration of 10 mM and was added to the plated cells as above. Cells were incubated at 37°C, 5% CO_2_ and 95% humidity for 24-72 hr. Using the Mosquito liquid handler, 300 nl of media was transferred from each well of two assay plates to respective white, solid bottom TC treated 1536-well recipient plates containing 4 μl/well 1x PBS every 24 hr. One media transfer plate received 3 μl/well NanoGlo substrate while the corresponding cell source plate received 4 μl/well coelenterazine-2P substrate. Luminescence was read on a ViewLux plate reader after a 5 min incubation. Coelenterazine-H was added at 4 μl/well to the second set of plates and luminescence was read after 5 min.

### qHTS Data Analysis

Assay statistics were calculated for 32 replicate wells of DMSO vehicle control and 2 wells of high concentration of respective compound treatment for initial assay optimization across the cell lines, cell densities, and for each substrate across a 72 hr time course. Statistics were then calculated for 64 wells of DMSO vehicle control and 32 replicate wells of max concentration of the respective control compounds for optimal assay conditions of 1500 cells/well treated for 48 hr. . Concentration response curves were determined for each compound and data was normalized to DMSO neutral control for each cell density for each cell line. Concentration response curves were fit and EC_50_ values calculated in Prism 7.0 (GraphPad, La Jolla, CA) with nonlinear regression log(agonist) vs. response – variable slope (four parameters) fit. Bell-shaped curves were also fit in Prism 7.0 with nonlinear regression and the equation:

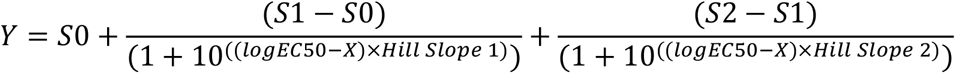

### 2-ACC Synthesis

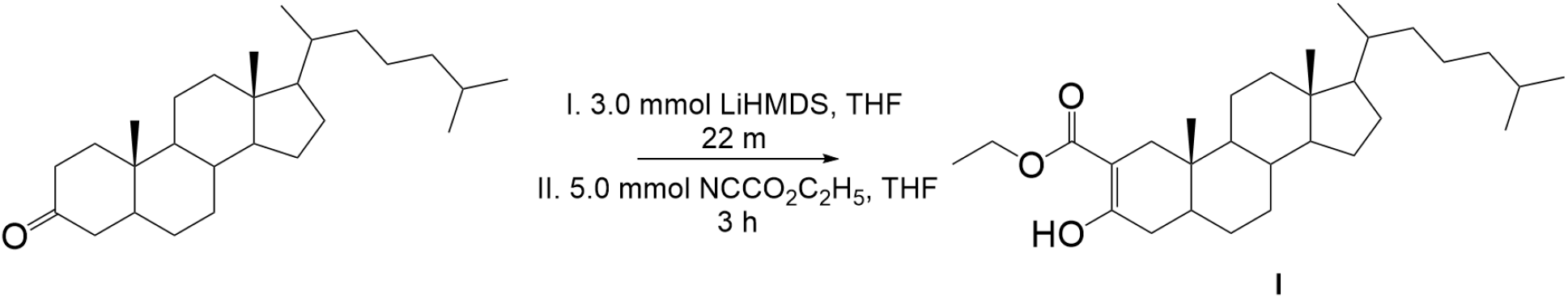

#### 2α-Ethoxycarbonylcholestan-3-one (I)

To a solution of cholestan-3-one (750 mg, 1.95 mmol) in dry THF (10 mL) under N_2_ at -78 °C was added 3 mL of LiHMDS (1M in hexane, 3.0 mmol) with stirring over 2 min. After another 20 min, ethyl cyanoformate (0.4 mL, 5.0 mmol) was added and the flask was removed from the bath and allowed to warm to room temperature. After 3 h the contents were poured into 150 mL of satd. aq. NaHCO_3_ and extracted three times with 40 mL of hexane / EtOAc 9:1. The combined organic phase was washed with 10% hydrochloric acid, brine, dried (Na_2_SO_4_) and evaporated under reduced pressure. Silica gel column chromatography (hexane / EtOAc 79:1) yielded the product as the enol tautomer (620 mg, 1.36 mmol, 70%). ^1^H NMR (600 MHz, CDCl_3_): 12.171 (1H, s), 4.26-4.15 (2H, m), 2.297 (1H, d, *J =* 15.6 Hz), 2.121 (1H, dd, *J* = 5.6, 18.8 Hz), 2.04-1.96 (2H, m), 1.82 (1H, m), 1.776 (1H, d, *J =* 15.6 Hz), 1.677 (1H, dq, *J* = 2.6, 12.9 Hz), 1.61-1.32 (m), 1.299 (3H, t, *J* = 7.1 Hz), 1.28-0.94 (m), 0.909 (3H, d, *J* = 6.4 Hz), 0.864 (6H, dd, *J* = 1.8, 6.5 Hz), 0.744 (3H, s), 0.668 (3H, s).

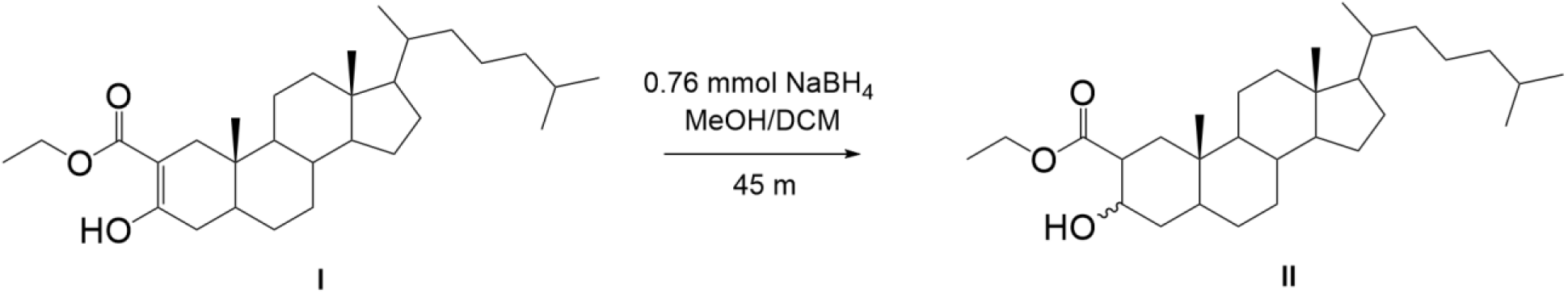

#### 2α-Ethoxycarbonylcholestan-3-ol (II)

To a solution of **I** (51.5 mg, 0.11 mmol) in MeOH / DCM 2:1 (3 mL) was added NaBH_4_ (29 mg, 0.76 mmol) in one portion. After 45 min, 10% hydrochloric acid (6 mL) was added and the mixture was extracted thoroughly with hexane / EtOAc 4:1. The extracts were filtered through silica gel and evaporated under reduced pressure to yield **II** (51.5 mg, 100%) as an inseparable mixture of stereoisomers (3β/3α = 2.2:1), distinguishable by the NMR shifts of the angular methyl groups and the coupling constants for H-3. ^1^H NMR (600 MHz, CDCl_3_), 3α: 4.206 (1H, tt, *J* = 2.3, 2.9 Hz), 0.816 (3H, s), 0.654 (3H, s). 3β: 3.592 (1H, tt, *J* = 4.9, 11.9 Hz), 0.715 (3H, s), 0.639 (3H, s).

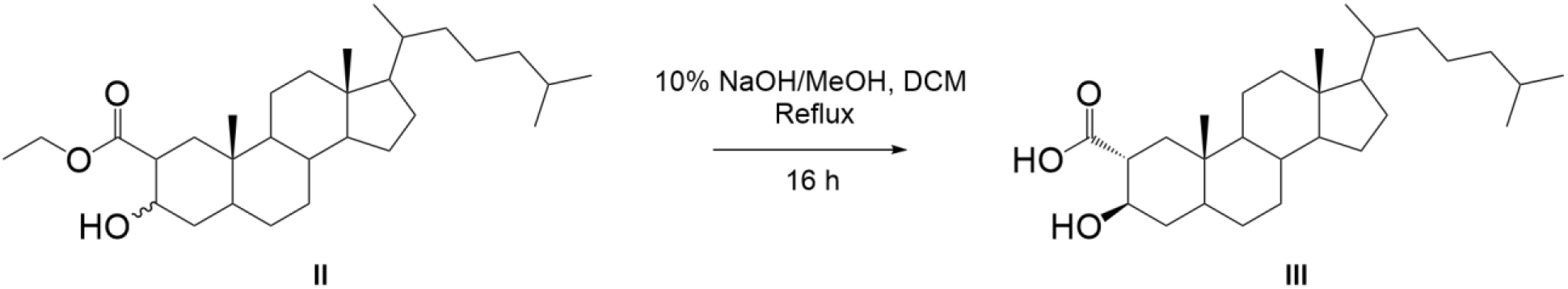

#### 3β-Hydroxycholestane-2α-carboxylic acid (III)

To a solution of **II** (22.0 mg mixed isomers, 0.048 mmol) in DCM (0.5 mL) was added 10% NaOH/MeOH (2.5 mL) and the solution was heated under reflux for 16 h. After cooling to rt, 10% hydrochloric acid (5 mL) was added, the mixture was extracted thoroughly with EtOAc, and the extracts were filtered through Na_2_SO_4_. Evaporation under reduced pressure yielded the product (20.6 mg, 100%). The diastereomers were separated by preparative TLC (250 μm silica plate, hexane / EtOAc 1:2 + 5% AcOH), and eluted with 2% AcOH/EtOAc. The more polar band contained the desired product. NMR (800 MHz, CDCl_3_): 3.818 (1H, td, *J* = 4.7, 10.6 Hz), 2.513 (1H, td, *J* = 3.0, 11.9 Hz), 2.048 (1H, dd, *J* = 3.2, 13.8 Hz), 1.965 (1H, dt, *J* = 3.0, 12.7 Hz), 1.809 (1H, m), 1.658 (2H, m), 1.58-1.47 (3H, m), 1.44-0.94 (m), 0.896 (3H, d, *J* = 6.4 Hz), 0.862 (6H, dd, *J* = 3.7, 6.6 Hz), 0.855 (3H, s), 0.673 (1H, td, *J* = 3.6, 11.4 Hz), 0.650 (3H, s). ^13^C NMR (201 MHz): 179.57, 71.14, 56.40, 56.23, 54.03, 46.79, 44.50, 42.55, 40.05, 39.87, 39.51, 36.16, 36.07, 35.78, 35.44, 31.84, 28.26, 28.21, 28.01, 24.18, 23.82, 22.81, 22.55, 21.30, 18.67, 12.52, 12.07. HRMS (HESI) *m/z* [M – H]^-^: Calcd for C_28_H_47_O_3_ 431.3525; Found 431.3532.

### CLZ-2-methylphosphonic acid (CLZ-P) synthesis

#### General Methods

All air or moisture-sensitive reactions were performed under a positive pressure of argon with oven-dried glassware. 4M HCl in dioxane and cesium carbonate were purchased from Sigma-Aldrich and used as such. Analytical analysis was performed on an Agilent LC/MS (Agilent Technologies, Santa Clara, CA). Method: A 7-minute gradient of 4% to 100% Acetonitrile (containing 0.025% trifluoroacetic acid) in water (containing 0.05% trifluoroacetic acid) was used with an 8-minute run time at a flow rate of 1 mL/min. A Phenomenex Luna C18 column (3-micron, 3 × 75 mm) was used at a temperature of 50 °C. A Phenomenex Gemini Phenyl column (3-micron, 3 × 100 mm) was used at a temperature of 50 °C. Purity determination was performed using an Agilent Diode Array Detector for both Method 1 and Method 2. Mass determination was performed using an Agilent 6130 mass spectrometer with electrospray ionization in the positive mode. ^1^H NMR spectra were recorded on Varian 400 MHz spectrometers. Chemical shifts are reported in ppm with undeuterated solvent (DMSO-*d*_6_ at 2.49 ppm) as an internal standard for DMSO-*d*_6_ solutions. High-resolution mass spectrometry was recorded on Agilent 6210 Time-of-Flight LC/MS system. Confirmation of molecular formula was accomplished using electrospray ionization in the positive mode with the Agilent Masshunter software (version B.02).

### Synthesis of diethyl ((4-(3,3-diethoxy-2-oxopropyl)phenoxy)methyl)phosphonate (3)

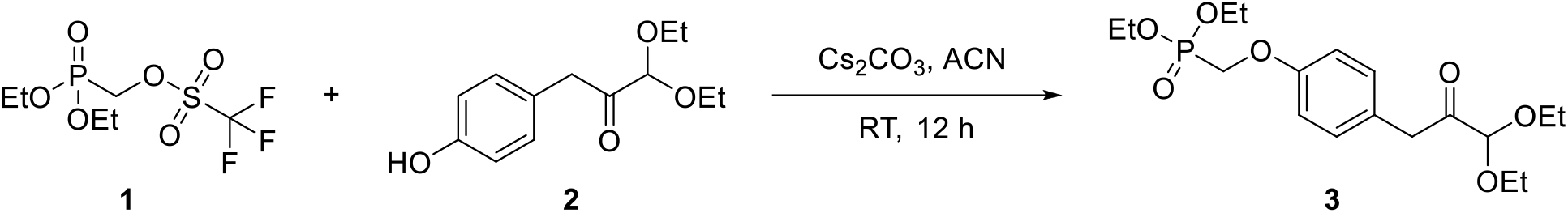

An oven-dried flask containing 1,1-diethoxy-3-(4-hydroxyphenyl)propan-2-one (0.794 g, 3.33 mmol, 1 eq) in acetonitrile (10 mL) was added cesium carbonate (1.628 g, 5.00 mmol, 1.5 eq). After stirring for 10 minutes at room temperature, (diethoxyphosphonyl)methyl trifluoromethanesulfonate (1 g, 3.33 mmol, 1 eq) was added and stirred at room temperature overnight. The reaction mixture was filtered through a pad of celite and washed with ethyl acetate. The filtrate was concentrated, and the residue was directly loaded to a flash 40 g silica column. The crude product was purified on a flash system eluting with 0-100 % ethyl acetate in hexanes over 32 column volumes to yield 0.62 g (48 %) pure product **3** after evaporating and drying process under high vacuum. LCMS Retention time = 5.3 min. (M+H)^+^ for C_18_H_30_O_7_P = 389.2.

### Synthesis of CLZ-2-methylphosphonic acid (CLZ-2P)

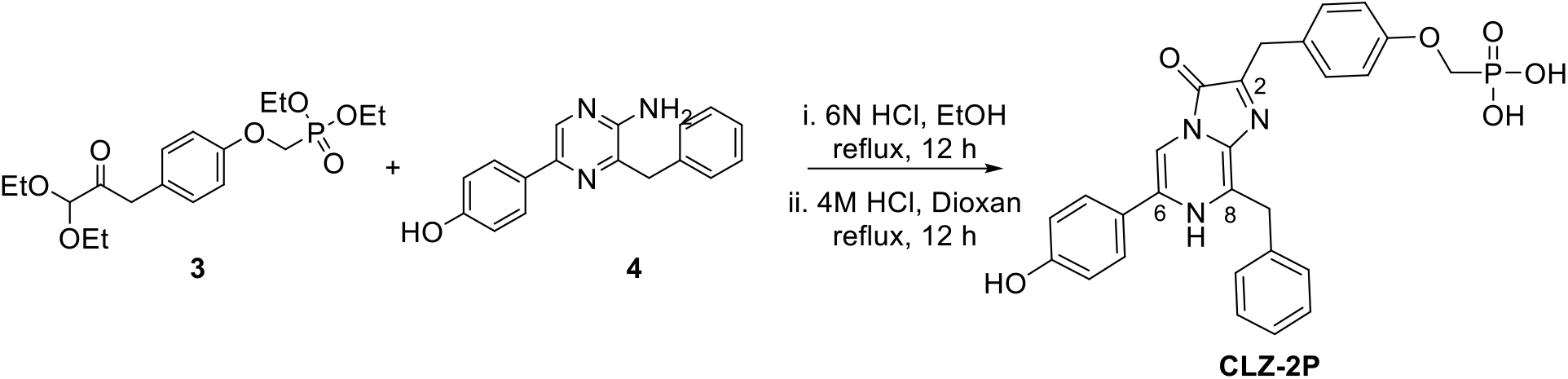

An oven-dried round-bottomed flask containing a mixture of diethyl ((4-(3,3-diethoxy-2-oxopropyl)phenoxy)methyl)phosphonate **3** (0.617 g, 1.589 mmol, 1.2 eq) and 3-benzyl-5-(4-(2- (trimethylsilyl)ethoxy)phenyl)pyrazin-2-amine **4** (0.5 g, 1.324 mmol, 1 eq; **4** was prepared following the literature method^48^) in degassed ethanol was added 6N HCl (in water) and refluxed overnight. The reaction was concentrated and added 100 mL of 4 M HCl in dioxan and refluxed again overnight at 100 °C to deprotect the ethyl phosphonate groups. The reaction was concentrated on a rotary evaporator and the crude product was purified on a reverse-phase flash system using 120 g C18 column eluting with 5-100 % acetonitrile (0.1 % TFA) in water (0.1% water). The product fraction was pooled and concentrated to obtain the relatively pure product which was further purified on an HPLC system eluting with water/acetonitrile (modified with 0.1 % NH_4_OH) using the basic method. A small portion was further subjected to HPLC purification using an acidic method to obtain pure material of yellow CLZ-2P as a TFA salt. LC-MS Retention Time = 3.54 min (M+H)^+^ for C_27_H_25_N_3_O_6_P = 518.1. ; 1H NMR (400 MHz, dmso) δ 10.90 (brs, 1H), 9.73 (brs, 1H), 7.82 – 7.52 (m, 2H), 7.44 (d, J = 7.5 Hz, 2H), 7.34 – 7.17 (m, 5H), 6.95 – 6.88 (m, 2H), 6.88 – 6.80 (m, 2H), 4.30 (s, 2H), 4.07 – 3.96 (m, 4H); HRMS (ESI) m/z (M+H)+ calcd. for C_27_H_25_N_3_O_6_P = 518.1475, found 518.1560.

