## Supplemental Information for "Bioluminescence-based reporters for characterizing inhibitors and activators of human Sonic Hedgehog protein autoprocessing in live cells at high throughput"

**PAGE 2.** **Supplementary Figure 1. 2-ACC activates *Drosophila* D46A Hh mutant protein in vitro**

**PAGE 3.** **Supplementary Figure 2. Amino acid sequence of HA-NLuc-SHhC reporter construct**

**PAGE 3.** **Supplementary Figure 3. Nucleic acid sequences used for CRISPR/Cas9 stable HA-NLuc-SHhC integration.**

**PAGE 4.** **Supplementary Table 1. Statistical characterization of D46A and WT reporter lines in 1536-well plate format**

**PAGE 5.** **NMR Spectra: 2-ACC**

**PAGE 7.** **NMR Spectra: CLZ-2P**

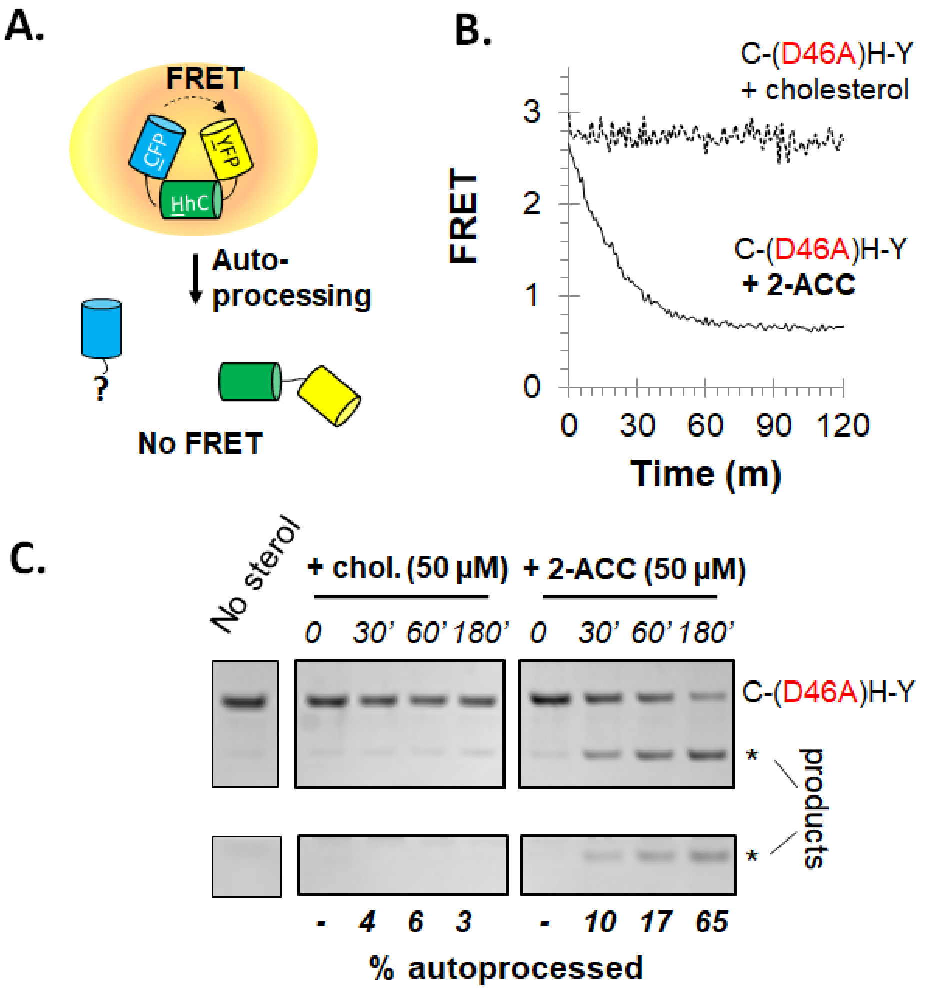


**Supplementary Figure 1.** 2-ACC activates *Drosophila* D46A Hh mutant protein in vitro. **A.** FRET-based optical reporter CFP-HhC-YFP (C-H-Y) detects Drosophila Hh autoprocessing by loss of FRET. **B**. FRET-active reporter mutant C-(D46A)H-Y, unreactive with cholesterol, readily accepts 2-ACC as a substrate. **C**) SDS-PAGE supports C-(D46A)H-Y autoprocessing with 2-ACC. Product fragments H-Y (52 kDa) and sterylated CFP (27 kDa).

METDTLLLWVLLLWVPGSTGD•YPYDVPDYAGAQPARSRDHMVLHEYVNAAGITSGSGAGVFTLEDFVGDWRQTAGY
NLDQVLEQGGVSSLFQNLGVSVTPIQRIVLSGENGLKIDIHVIIPYEGLSGDQMGQIEKIFKVVYPVDDHHFKVILHYGTLVI
DGVTPNMIDYFGRPYEGIAVFDGKKITVTGTLWNGNKIIDERLINPDGSLLFRVTINGVTGWRLCERILATGLQKAENSVAA
KSGG▼**(C)**FPGSATVHLEQGGTKLVKDLSPGDRVLAADDQGRLLYSDFLTFL**(D)**RDDGAKKVFYVIETREPRERLLLTAA
HLLFVAPHNDSATGEPEASSGSGPPSGGALGPRALFASRVRPGQRVYVVAERDGDRRLLPAAVHSVTLSEEAAGAYAPL
TAQGTILINRVLASCYAVIEEHSWAHRAFAPFRLAHALLAALAPARTDRGGDSGGGDRGGGGRVALTAPGAADAPGAG
ATAGIHWYSQLLYQIGTWLLDSEALHPLGMAVKSS

YPYDVPDYAGAQPARSRDHMVLHEYVNAAGITSGSGAGVFTLEDFVGDWRQTAGYNLDQVLEQGGVSSLFQNLGVSV
TPIQRIVLSGENGLKIDIHVIIPYEGLSGDQMGQIEKIFKVVYPVDDHHFKVILHYGTLVIDGVTPNMIDYFGRPYEGIAVFD
GKKITVTGTLWNGNKIIDERLINPDGSLLFRVTINGVTGWRLCERILATGLQKAENSVAAKSGG-sterol

**Supplementary Figure 2**. Amino acid sequence of NLuc-SHhC reporter construct. *Top*. Precursor HA-NLuc-SHhC. Secretion signal peptide (purple); HA-tag (yellow); NLuc (cyan); SHhC (grey). Catalytic residues, C1 and D46, of SHhC, shown in bold, are mutated to alanine in the **C1A** and **D46A** reporter constructs. *Bottom*. HA-Nluc-sterol product expected from SHhC autoprocessing, with sterol esterified to the C-terminal glycine.

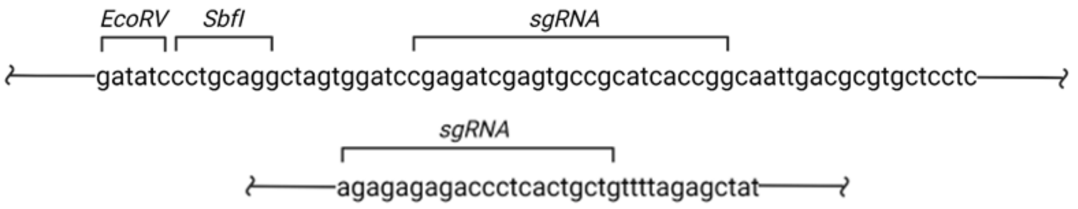


**Supplementary Figure 3**. Nucleic acid fragments used for CRISPR/Cas9 facilitated HA-NLuc-SHhC integration. *Top*. CRISPR targeting sequence inserted into SnaBI site of pDisplay vector. *Bottom*. Short guide RNA sequence used for targeted integration of **WT** and **D46A** reporter plasmids.

| Parameters | Coelenterazine-2P NLuc Substrate | | | | CellTiter-Glo Cytotoxicity Counter-Screen | | | |
|---|---|---|---|---|---|---|---|---|
| | D46A | | WT | | D46A | | WT | |
| S/B | 1.8 | | 9.6 | | 21.3 | | 20.2 | |
| Number of Wells | DMSO 64 | 2-ACC 32 | DMSO 64 | CHX 32 | DMSO 64 | Digitonin 32 | DMSO 64 | Digitonin 32 |
| Output Signal | DMSO 27.6 ± 1.2 | 2-ACC 50.8 ± 1.6 | DMSO 6,255.9 ± 310.7 | CHX 652.9 ± 25.2 | DMSO 13,541.5 ± 474.8 | Digitonin 637.2 ± 95.5 | DMSO 16,435.9 ± 597.0 | Digitonin 812.5 ± 146.3 |
| CV | 4.5 | 3.1 | 5.0 | 3.9 | 3.5 | 15.0 | 3.6 | 18.0 |
| Z' factor (Mean) | 0.64 | | 0.83 | | 0.87 | | 0.86 | |
| Control condition 1 | 2α-Carboxy Sterol (28.75 µM) | | 57.5 µM Cycloheximide | | Digitonin (115 µM) | | Digitonin (115 µM) | |

**Supplementary Table 1**. Statistical characterization of miniaturized **D46A** and **WT** reporter lines in 1536-well plate format treated with CLZ-2P. NLuc-SHhC assay assessed for media bioluminescence where **D46A** mutant samples were normalized to 28.75 µM 2-ACC as 100% stimulation control and **WT** cells were assessed for inhibition and normalized to 57.5 µM CHX control as 100% inhibition. CellTiter-Glo cytotoxicity assay assessed for inhibition of luminescence where both cell lines were normalized to 115 µM Digitonin as 100% cytotoxicity control. Statistics were calculated from one 1536-well plate with 32-64 replicate wells of respective controls.

### NMR spectra for 2-ACC synthesis

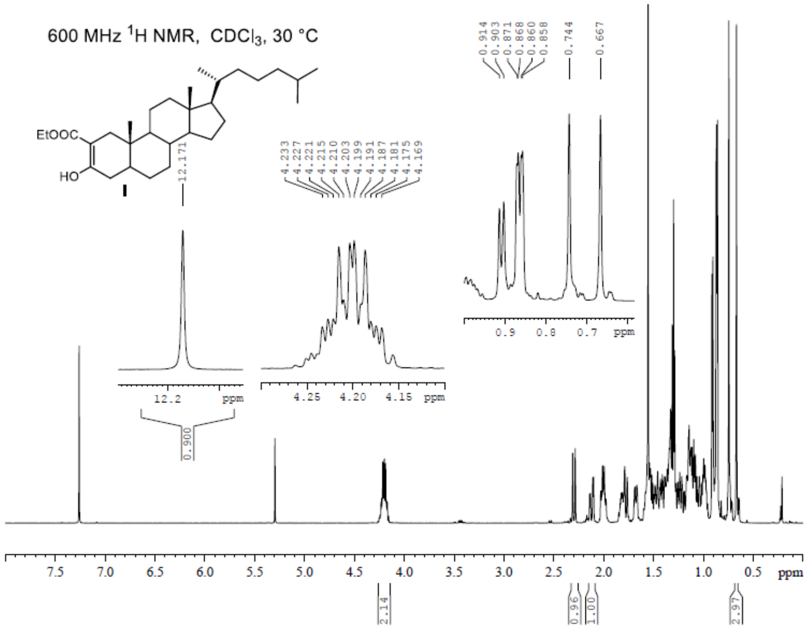


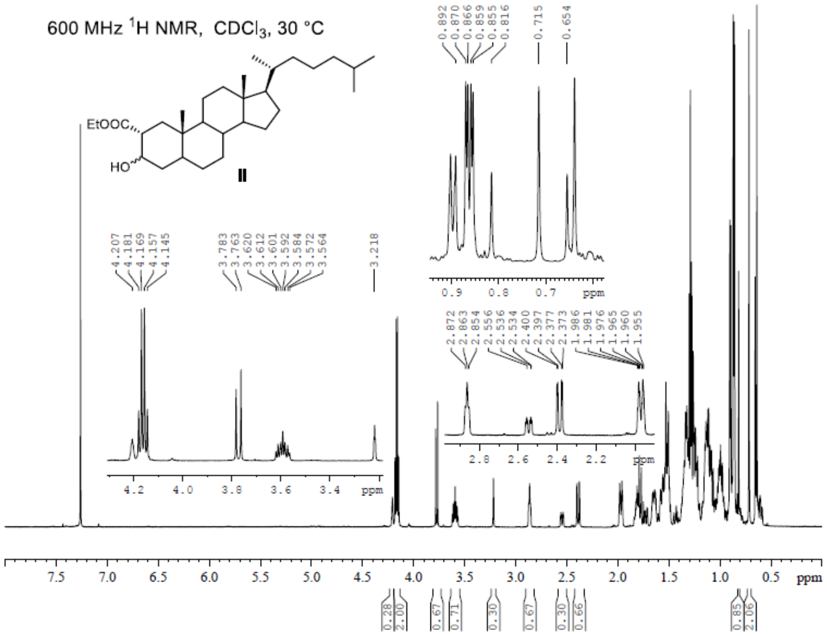

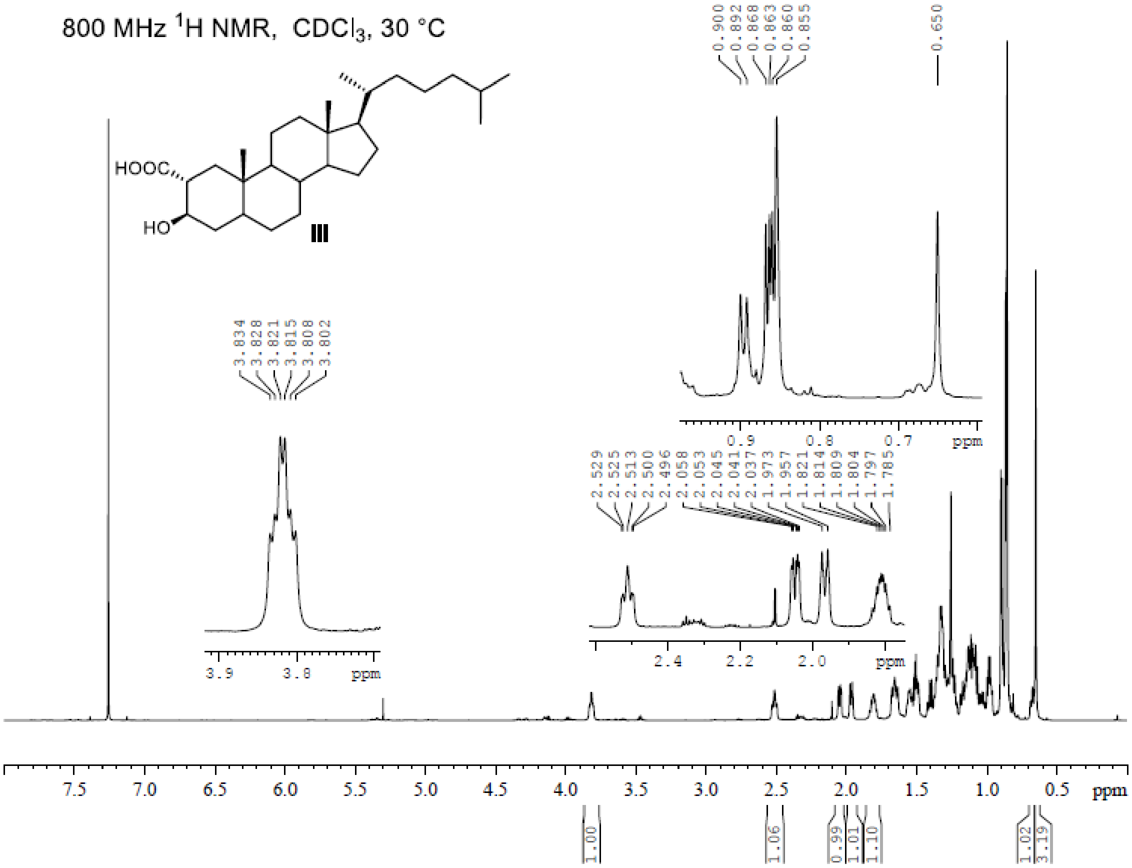
800 MHz ¹H NMR, CDCl₃, 30 °C
HOOC
HO
III
ppm

### NMR Spectra: CLZ-2P

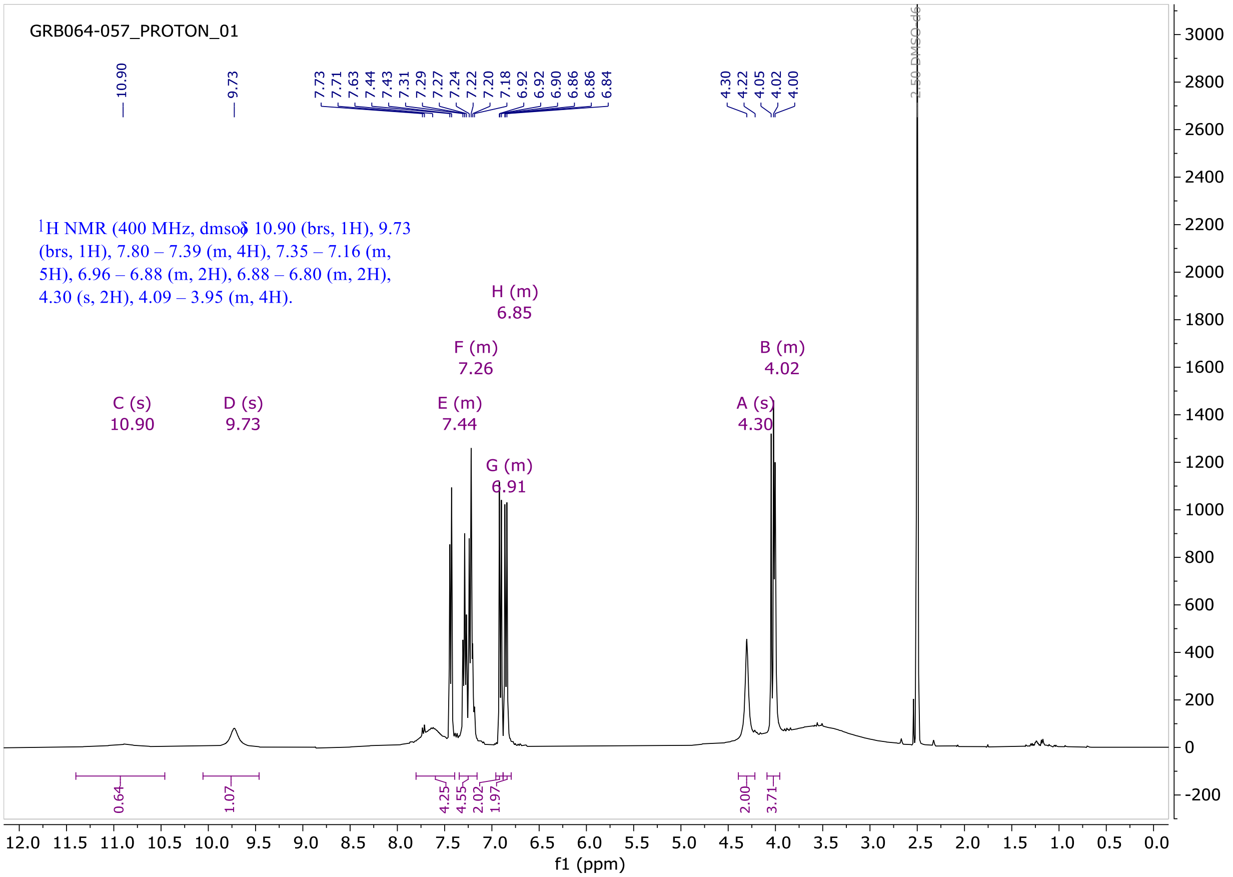
